# Genome-wide association study of carbon and nitrogen metabolism in the maize nested association mapping population

**DOI:** 10.1101/010785

**Authors:** Nengyi Zhang, Yves Gibon, Nicholas Lepak, Pinghua Li, Lauren Dedow, Charles Chen, Yoon-Sup So, Jason Wallace, Karl Kremling, Peter Bradbury, Thomas Brutnell, Mark Stitt, Edward Buckler

**Author notes:** Present address: BASF Plant Science, 26 Davis Dr., Research Triangle Park, North Carolina 27709, USA. Present address: INRA, UMR 1332, Univ.Bordeaux, F-33883 Villenave d’Ornon, France. Present address: Department of Crop Science, Chungbuk National University, Cheongju, South Korea (Yoon-Sup So). These authors contributed equally to this work (Nengyi Zhang, Yves Gibon).

## Abstract

Carbon (C) and nitrogen (N) metabolism are critical to plant growth and development and at the basis of yield and adaptation. We have applied high throughput metabolite analyses to over 12,000 diverse field grown samples from the maize nested association mapping population. This allowed us to identify natural variation controlling the levels of twelve key C and N metabolites, often with single gene resolution. In addition to expected genes like *invertase*s, critical natural variation was identified in key C_4_ metabolism genes like *carbonic anhydrase*s and a *malate transporter*. Unlike prior maize studies, extensive pleiotropy was found for C and N metabolites. This integration of field-derived metabolite data with powerful mapping and genomics resources allows dissection of key metabolic pathways, providing avenues for future genetic improvement.

---

Carbon (C) and nitrogen (N) metabolism are the basis for life on Earth. The production, balance and tradeoffs of C and N metabolism are critical to all plant growth, yield, and local adaptation (*1, 2*). In plants there is critical balance between the tissues that are producing energy (sources) and those using it (sinks), as this change varies through time and developmental stage (*3*). While a great deal of research has focused on the key genes and proteins involved in these process (*4-6*), relatively little is known about the natural variation present within a species that fine tune these processes.

In the last two decades, quantitative trait loci (QTL) mapping, first with linkage analysis, later with association mapping, has been used to dissect C and N metabolism in Arabidopsis (*7-10*), tomato (*11*) and maize (*12-16*). Because of the drawbacks of both linkage analysis and association mapping, these studies are of either low resolution or low power. Additionally, most of these studies have not focused on C and N metabolism in the field, where adaptation is most relevant. We used a massive and diverse germplasm resource (the maize nested association mapping (NAM) population) that captures representative diverse alleles from around the world (*17, 18*) to evaluate genetic variation of 12 metabolites in a maize field (Fig. 1). These metabolites are key indicators of photosynthesis, respiration, glycolysis, and protein and sugar metabolism in the plant (*9*). The 5,000 lines of NAM are the product of crossing 25 diverse lines to a reference line, B73, whose genome sequence has been published (*19*). Importantly, this reference design permitted all the germplasm to be grown and evaluated in the same field for metabolic profiling. Additionally, the NAM population permits gene-level resolution of complex traits, even in the field (*20-22*). By taking advantage of a robotized metabolic phenotyping platform (*23*), we were able to conduct a large study of plant metabolism, with more than 100,000 assays conducted across over 12,000 samples (2 independent samples per experimental plot).

**Fig. 1.**
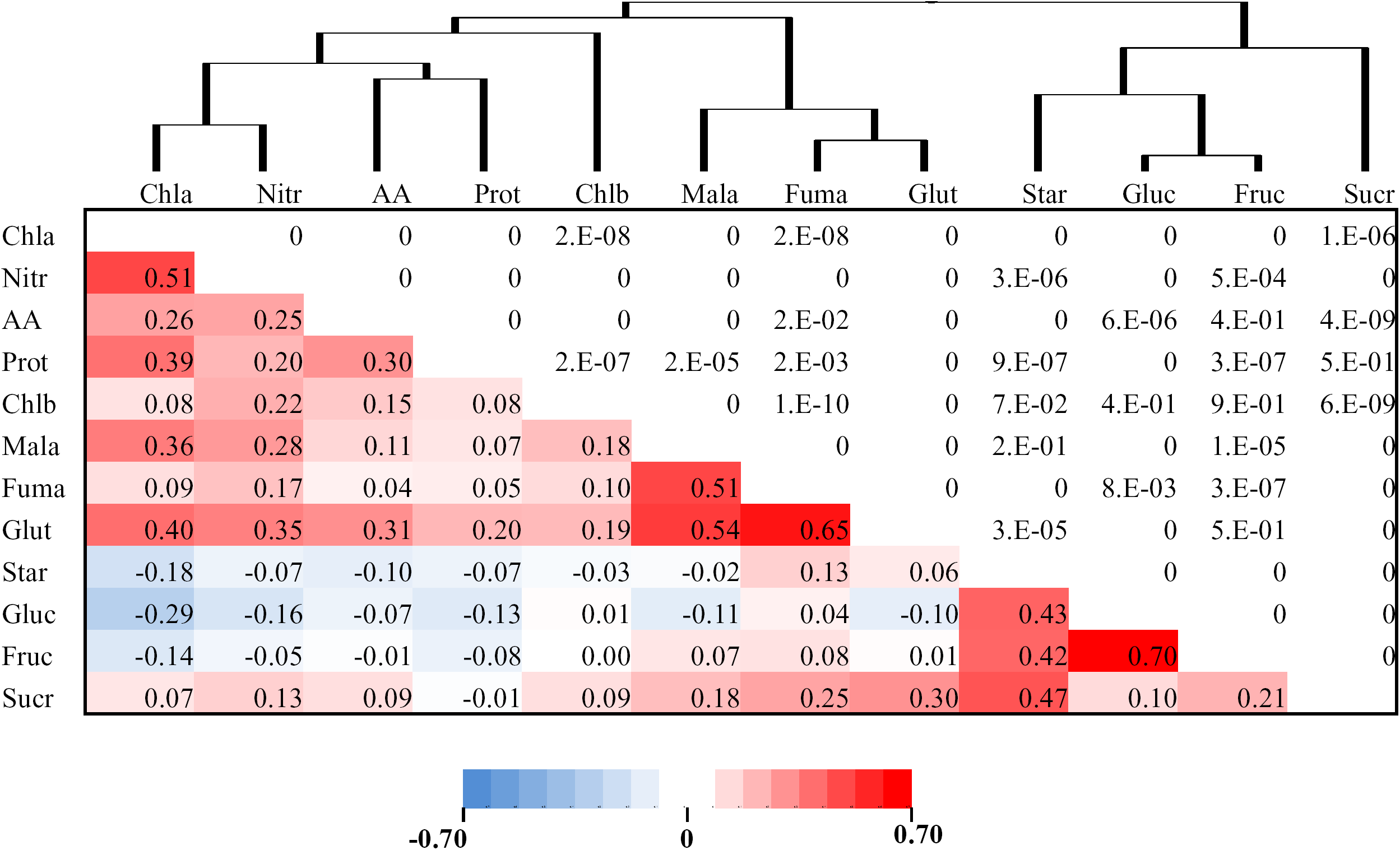
Correlation matrix and clusters of the 12 carbon and nitrogen metabolites. Lower left part of the matrix is correlation coefficients accounted for the maturity effect and upper right part is the corresponding *p*-values. *p*-value < 1E-10 was rounded down to 0. *p*-value < 8E-4 is significant at a Bonferroni correction at α = 0.05. Chla, chlorophyll a; Nitr, nitrate; AA, total amino acids; Prot, protein; Chlb, chlorophyll b; Mala, malate; Fuma, fumarate; Glut, glutamate; Star, starch; Gluc, glucose; Fruc, fructose; Sucr, sucrose.

A complication conducting diversity studies of this scale is that sampling has to be performed over a period of time relative to flowering to approximate similar developmental time points. However, while we could not control for weather or development rate experimentally, with the use of such a large sample, we were able to statistically remove the effects of weather, day and time of sampling, small scale spatial effects, and differences in maturity. For the 12 key metabolites, whole field repeatabilities were between 15% and 71% and broad-sense heritabilities (*H*^2^) across the NAM founder lines were between 14% and 68% (table S1), indicating the high potential for accurate mapping of the genes responsible for the metabolic variation.

For most of the developmental and disease traits studied so far in maize (*17, 20-22*) there is a strong correlation between traits across diverse germplasm panels but careful studies of pleiotropy in NAM has suggested few or no QTL are shared (*20, 22*). Therefore, this kind of phenotypic correlation is due to stacking of different genes each controlling an independent trait through evolution and breeding, rather than due to variation in shared genetic mechanisms. The metabolites analyzed in the present study showed substantial correlations across the NAM population and within each subpopulation. Correlations and clustering were observed, as expected, between metabolites within N portions of the pathways (Fig. 1), and within the starch-sugar pathways, respectively. Some traits like chlorophyll b (Chlb) did not substantially correlate with other traits. As we show below in joint linkage analysis (Fig. 2A and fig. S1) and genome-wide association studies (GWAS) (Figs. 3A and 3B, and tables S2 and S3) there is substantial overlap in QTL between metabolic traits. Further, metabolites that show correlated changes across the entire NAM population (Fig. 1) had a significantly higher frequency of shared QTL (fig. S2. *R*^2^=0.83, *p* =2.1E-26). This suggests that, in contrast to most other traits studied to date in maize, there are a relatively small number of control points for the natural variation despite the complexity of N and C metabolism.

**Fig. 2.**
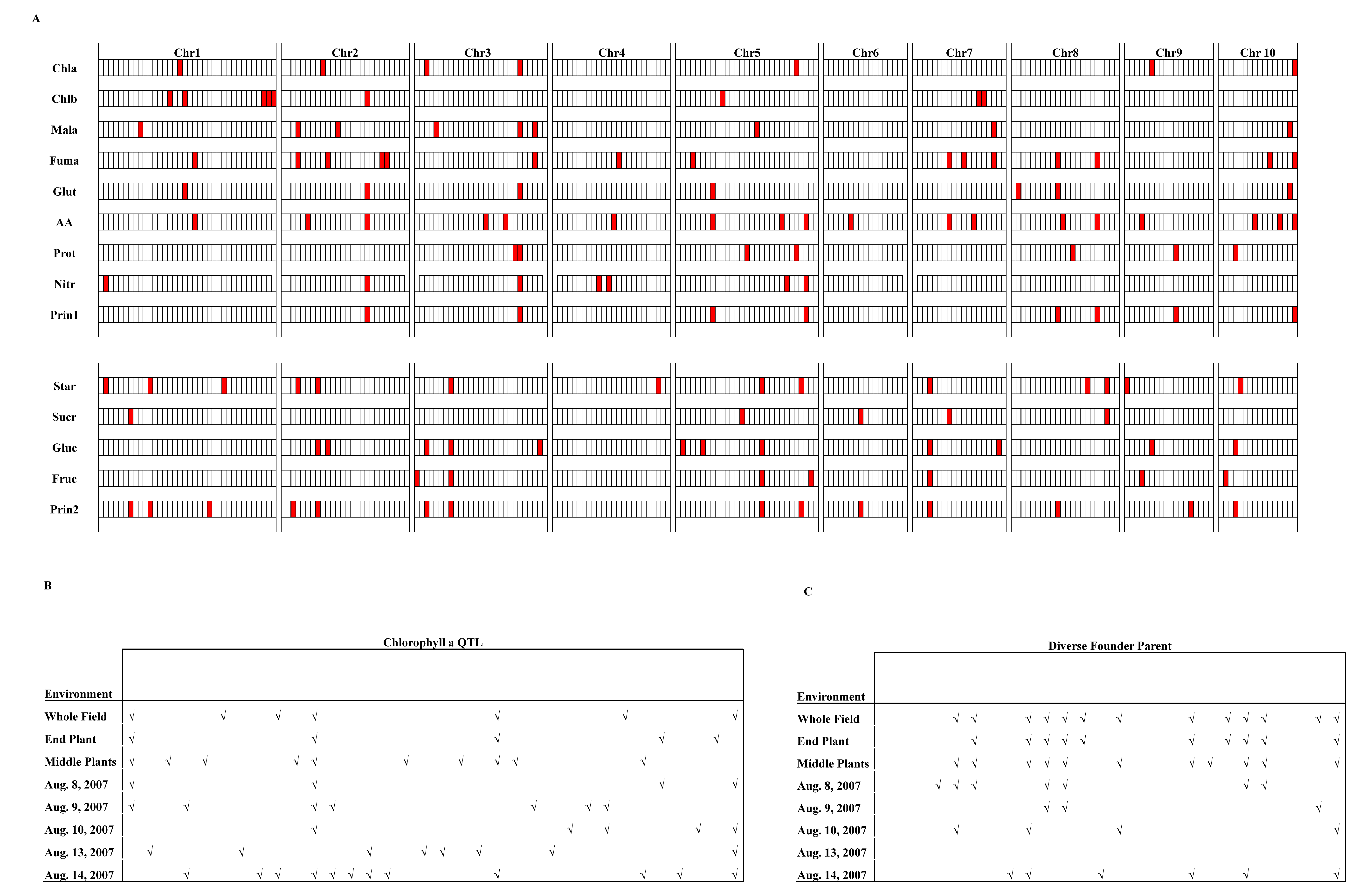
QTL distribution across different chromosomes. (A) QTL distribution of the 12 carbon and nitrogen metabolites and the first two principal components derived from the whole field. (B) QTL distribution of Chla derived from different environments. (C) QTL m411distribution in different NAM families in different environments. Mala, malate; Fuma, fumarate; Glut, glutamate; Chla, chlorophyll a; Chlb, chlorophyll b; Nitr, nitrate; Sucr, sucrose; Gluc, glucose; Fruc, fructose; Star, starch; AA, total amino acids; Prot, protein; Prin1, first principal component; Prin2, second principal component. The analysis was based on data derived from the whole field, the end plant in a row, the middle plants in a row, and different day of sampling.

**Fig. 3.**
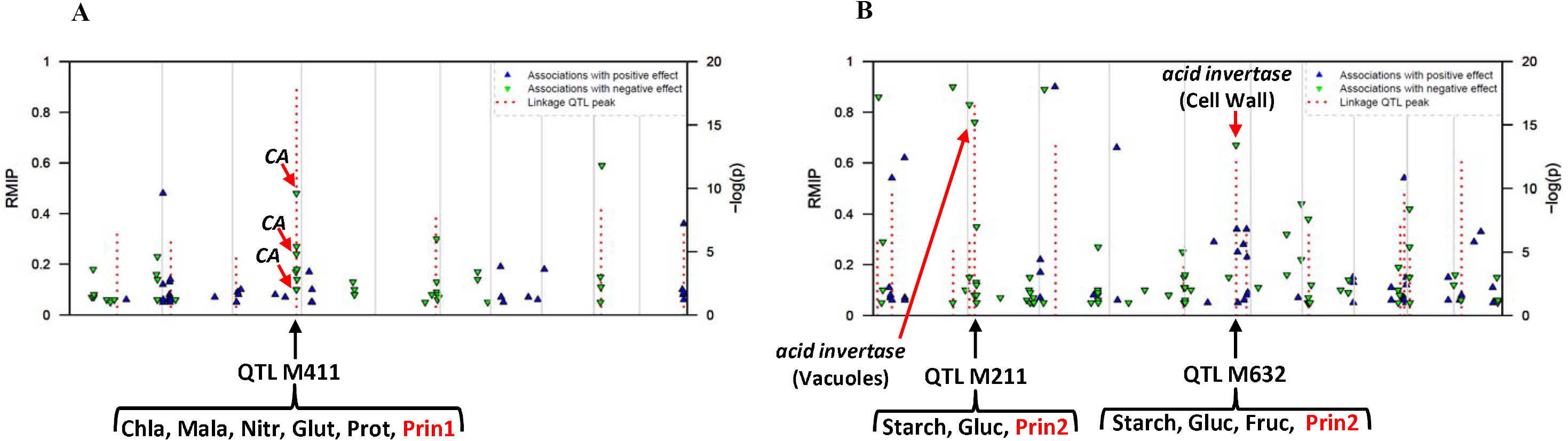
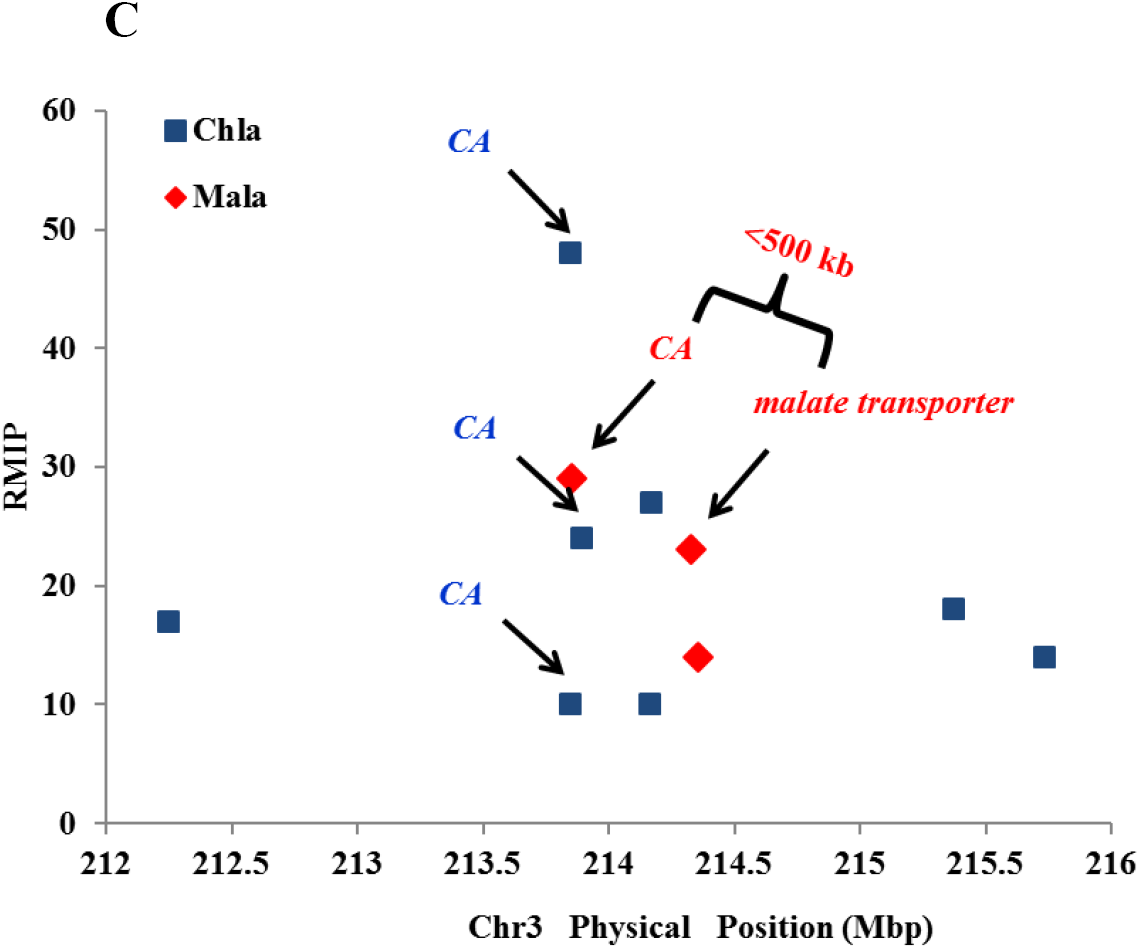
Associations between metabolites and SNPs. All SNPs detected as significant in at least 5 subsamples are triangles (blue with positive effect and green with negative effect) relative to their physical sequence position. Vertical positions of triangles represent resample model inclusion probability (RMIP) of the SNP. QTL are red lines whose vertical positions represent their *F*-test log(1/*P*) in the final joint linkage QTL model. **(A)** Chla and SNPs from *carbonic anhydrases* (*CAs*) on Chromosome 3 (Chr3). *CA* region on Chr3 overlaps with QTL M411. Chla, Mala, Glut, Nitr, Prot, and Prin1 share the same QTL M411 and, moreover, they all have significantly associated SNPs from the *CA*s in this QTL. Chla, chlorophyll a; Mala, malate; Nitr, nitrate; Glut, glutamate; Prot, protein; Prin1, first principal component. **(B)** Star and SNPs from *invertases* on chromosomes 2 (Chr2) and 5 (Chr5)*. invertase* region on Chr2 overlaps with QTL M211, which Star, Glu, and Prin2 share, and on Chr5 overlaps with M632, which Star, Glu, Fru, and Prin2 share. Gluc, glucose; Fruc, fructose; Star, starch; Prin2, second principal component. **(C)** Associations identified by GWAS at *carbonic anhydrases* (*CA*) and a *malate transporter* region on chromosome 3 (Chr3). Vertical positions of diamonds represent resample model inclusion probability (RMIP) of the SNP. Chla, chlorophyll a; Mala, malate.

Clustering and principal component analysis (PCA) of traits effectively summarized the variation in these pathways (Fig. 1 and fig. S3). These summary statistics were one focus of the GWAS analysis. Since there were relatively few axes of variation, we summarized the data with PCA after accounting for the flowering (days to anthesis, DTA) effect (*24*). The first two principal components (PCs; Prin1 and Prin2) explained 25% and 20% of the variation (fig. S3A). In Prin1, N metabolites showed positive and higher contributions than C metabolites while the opposite appeared in Prin2 (fig. S3B).

Using joint linkage analysis (*17*), we identified 5-18 QTL for the 12 metabolites and the first two PCs explaining 24%-55% of the phenotypic variation (Fig. 2A tables S1 and S4). We included DTA in the model for the mapping of each metabolite. Many QTL have both negative and positive effect alleles relative to B73 among the NAM founder lines (fig. S4). On average, each QTL is shared by six families (fig. S5). We did not detect digenic epistatic interactions between QTL with additive effects. Small effect QTL were detected for all 12 metabolites and for the first two PCs. Most QTL explain 2% or less of the variance (fig. S6) and most QTL alleles were estimated to give a less than 2% of increase or decrease change (fig. S7). Similar results have been observed for whole plant traits (*17, 20, 25*) and diseases resistance traits (*21, 22*) in maize. These results suggest that primary metabolic traits have a similar genetic architecture to complex traits in maize.

To identify the genes controlling this variation, we applied the recently published GWAS method (*20*) to test the association of 1.6 million maize HapMap SNPs (*26*). The resample model inclusion probability (RMIP) was calculated based on how often each SNP is included in each model generated from the resampled data (*27, 28*). RMIP identified 1,394 significantly associated SNPs (RMIP ≥ 0.05; Figs. 3A and 3B, fig. S8, and tables S2 and S3). Predicted genes containing, or directly adjacent to, SNP associations were evaluated as potential candidate genes for metabolites. Interestingly, we identified many genes which are known to be involved in plant C and N metabolic pathways (Figs. 3A and 3B and table S2).

We detected three SNPs from *carbonic anhydrases* (*CA*s) on chromosome 3 (Chr3) that are associated with chlorophyll a (Chla) variation (Fig. 3A and table S2). There are 5 paralogs of *CA* in B73 genome and, among them, two tandemly duplicated on Chr3. Two of the three significant SNPs are from the one paralog (*CA3.1*) and the other from the other paralog *(CA3.2)* on Chr.3. The Chr3 *CA* region overlaps with QTL M411. This QTL is very stable across different environments, such as it existed in different part of the field (End Plant and Middle Plants) and four of the five sampling days in the same or different families (Figs. 2B and 2C). The N metabolites, Chla, malate, nitrate, glutamate and protein as well as Prin1 share the same QTL and, moreover, all have significantly associated SNPs from the *CA*s in this QTL (Fig. 3A and table S2). Moreover, single gene resolution has been achiveved in this region of the NAM population (fig. S9A). The cytosolic isoform of CA catalyzes the first step in the C_4_ photosynthetic pathway in mesophyll (M) cells, resulting in the hydration of CO_2_ to bicarbonate (*29*). Bicarbonate is then fixed by phosphoenolpyruvate carboxylase to produce oxaloacetate (OAA). OAA is then reduced to malate, which diffuses via plasmodesmata into the bundle sheath (BS) cells (*30*) where it is decarboxylated by an NADP-dependent malic enzyme (NADP-ME) to generate a high internal CO_2_ concentration in BS plastids. There exists another minor decarboxylation system with phospho*enol*pyruvate carboxykinase (PEPCK) in addition to the major one with NADP-ME in maize. This C_4_ carbon shuttle, results in elevated CO_2_ concentration in the BS nearly completely suppressing the energy wasteful process of photorespiration (*31*). Among the significant *CA* SNP associated metabolites, malate is a key metabolic intermediate in C_4_ pathway, total protein is a central metabolic and cellular parameter and will, among other things, affect photosynthetic capacity, Chla is related to light-dependent photosynthesis reactions, nitrate is the major source of N for proteins and chlorophylls, and glutamate is the product of nitrate and ammonium assimilation and the precursor for the biosynthesis of amino acids and many other N-metabolites including Chla and Chlb. Moreover, Prin1, which captured the common factor of N metabolites, is also significantly associated to SNPs from *CA*s on Chr.3. Therefore, genetic variation of *CA*s on Chr3 is associated with N metabolism and photosynthesis. In Arabidopsis, CAs are upstream regulators of CO_2_-controlled stomatal movements in guard cells so they can have significant implications in plant water use efficiency (*32*). In C_4_ grasses CA levels are thought to be rate limiting for photosynthesis (*33, 34*)

In B73, most of the *CA* expression in seedlings is derived from the two paralogs *CA3.1* and *CA3.2* on Chr.3 (fig. S10) (*35*) which is consistent with our QTL and GWAS results, though the expression experiments were conducted at 3-leaf stage instead of flowering time (*35*). Thus, we performed real-time PCR on the two Chr.3 *CA* paralogs in the NAM founders using material sampled at flowering time, the same developmental stage as the QTL and GWAS experiment. The result showed that transcript for the B73 allele is significantly higher than the alternative allele for the first paralog *CA3.1* and marginally higher for the second paralog *CA3.2* (Fig. 4A). However, the variance of the expression for the first paralog is also significantly different between the two classes of alleles (Fig. 4B), which indicates that we might not capture the causal SNP which is responsible for the metabolite variation.

**Fig. 4.**
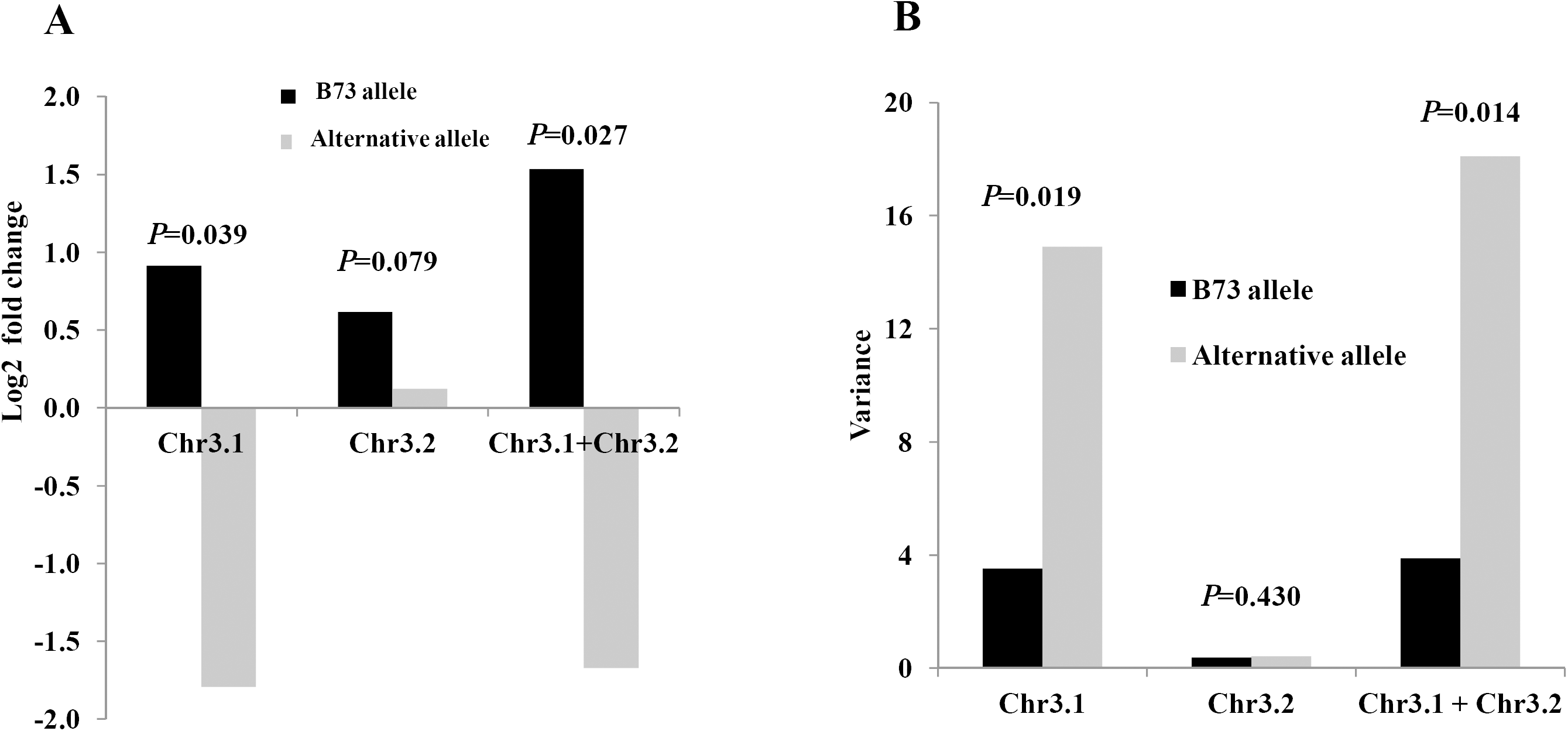
Expression and corresponding variance comparison for *carbonic anhydrases* between B73 allele and alternative allele in NAM founder lines at flowering time. **(A)** Expression. **(B)** Variance.

For malate, we also identified a significantly associated SNP adjacent to a malate transporter gene (~2 kb) which is about 500 kb away from the significantly associated *CA* SNP in QTL M411 region (table S2 and Fig. 3C). Malate transporters are required for the movement of malate between mitochondria, cytosol, vacuole, and chloroplast in both M and BS cells and, therefore, it is also important to photosynthesis (*36-38*).

We detected a significant SNP (RMIP=0.76) from a vacuolar *acid invertase* in the major QTL M211 region and a SNP (RMIP=0.67) from a cell wall *acid invertase* in the QTL M632 region associated with starch variation (Fig. 3B). Starch, glucose, and Prin2 share the QTL M211 and starch, glucose, fructose and Prin2 share the QTL M632. It is well known that invertases regulate C metabolism (*39-41*).

We identified several other candidate genes with recognizable roles in C and N metabolism (table S2). For example, we identified a *starch synthase* relevant to Chla variation, a *trehalose-6-phosphate synthase* and a *nitrate transporter* relevant to glucose variation, and two *cellulases* related to fructose and glucose, respectively. We also detected a ribosomal protein associated to protein content, which might be of great interest as protein synthesis is intimately linked to biomass production (*42*). Three genes assigned to cell wall synthesis and modification are found among genes that are associated with protein content. Protein content is likely to be determined not only by protein synthesis per se but also by the rate of cell expansion, which will be affected by, among other process, cell wall synthesis and modification. We identified Chla,b binding protein, glutamine synthetase, NADP-malate dehydrogenase and PEPC kinase as associated to nitrate. Glutamine synthetase is involved in the nitrate and ammonium, NADP-malate dehydrogenase is involved in the export of reducing equivalents from the chloroplast (*43*), to support reductive reactions (like nitrate reductase) in the cytosol and the formation of ATP in mitochondria and is a central component to the C_4_ carbon shuttle in M cells (reducing OAA to malate). PEPC kinase regulates the activity of PEP carboxylase, which as mentioned above is a component of the C_4_ carbon shuttle, and is also involved in the synthesis of malate that acts as a counteranion during nitrate assimilation (*44*). We also identified pyruvate dehydrogenase E1 as relevant to fumarate variation.

We observed significant enrichment of association around candidate genes for C and N metabolites. Out of 514 candidate genes potentially related to C and N metabolism curated from maize pathway analyses (table S5), 101 are overrepresented with GWAS signals (fig. S11). Those significantly enriched candidates include above mentioned *CA*s, *invertase*s and the *malate transporter*. Through GO term enrichment analysis, most of the *a priori* candidates discovered in our GWAS take part in carbohydrate metabolic, alcohol metabolic, carbohydrate catabolic, and glucose metabolic processes (table S6). We also compared the distribution of different classes of genes which the 126 most significantly associated SNPs (RMIP>=0.50) reside in or are adjacent to with that in the whole maize genome. The results showed that the C_4_ and source-sink genes are over represented (fig. S12). However, most SNPs are still located in or adjacent to genes from which we do not know yet if and how they are related to C and N metabolism (table S3).

Currently, C_4_ grasses are some of the most productive plants on Earth (*45*). However, many of our key crops such as rice, wheat, and potato perform C_3_ photosynthesis and a major international effort is underway to convert some of these crops to C_4_ (*46*). This study clearly shows that fine tuning of the expression of genes that drive the C_4_ carbon shuttle are likely key determinants of local adaptation and hence yield. Our results show that levels of central metabolites in C and N metabolism are determined by genetic variation in key genes involved in CO_2_ capture (carbonic anhydrase) and movement (malate transporters). These proteins likely impact sub-processes in C_4_ photosynthesis, including the extent to which the C_4_ cycle draws down CO_2_ in the M airspaces, and the movement of malate from the M into the BS. Manipulation of these loci by breeding or transformation and then optimization in a wide range of environments could lead to even more efficient yield. Despite the variation present today, it is unlikely that a tropical grass has evolved all the appropriate alleles to fix carbon well not only in the cold temperate springs, but also in the hot summer. Thus, these high resolution diversity studies can help guide breeding efforts to identify genes that are most likely to be amenable to genetic improvement.

## Acknowledgments

We thank S. Miller for technical editing of the manuscript; researchers in Buckler Laboratory at Cornell University for sample collection and data analyses, researchers in Stitt Laboratory at Max Planck Institute of Molecular Plant Physiology for assistance with the metabolite measurements, and Marcela Monaco in Ware Laboratory at Cold Spring Harbor Laboratories for providing some of the candidate genes. Mention of trade names or commercial products was solely to provide specific information and does not imply recommendation or endorsement by the USDA. This work was supported by NSF grants DBI–0501700, DBI–0321467, and DBI–0820619, the USDA-ARS and Max Planck Society.

